# Comparing Machine Learning Architectures for the Prediction of Peptide Collisional Cross Section

**DOI:** 10.1101/2022.03.01.482566

**Authors:** Emily Franklin, Hannes L. Röst

**Affiliations:** University of Toronto

## Abstract

Mass spectrometry is the method of choice in large-scale proteomics studies. One common method is data-independent acquisition (DIA), which allows for high-throughput analysis of biological samples, but also produces complex data. Methods of peptide separation, in addition to retention time, improve data analysis and there has been increasing interest in separating peptides based on collisional cross section (CCS), which is a measure of the size of a peptide. However, existing libraries that are used during data analysis lack CCS measurements, and this data is expensive and time-consuming to acquire. This has led to the desire to predict library values for mass spectrometry analysis. Here we compare three deep learning architectures, LSTM, CNN, and transformer, for the tasks of retention time and collisional cross section prediction. We show that the LSTM and CNN models perform similarly and that the transformer has a lower performance than expected.

## 2 Introduction

Large scale proteomics is playing an increasingly important role in medicine, environmental science, forestry and other industries where studying the entire collection of proteins present in an organism or tissue can produce useful biological insights. Liquid chromatography coupled with tandem mass spectrometry (LC-MS/MS) is a widely used technique for proteome analysis due to its high throughput compared with traditional techniques.

There are two main approaches for mass spectrometry analysis: data-dependent acquisition (DDA) and data-independent acquisition (DIA). Both are bottom-up proteomics approaches and follow the usual pattern of protein digestion into peptides with protease enzymes followed by separation via liquid chromatography then mass spectrometry. In DDA, peptides are chosen online for fragmentation prior to a second round of mass spectrometry analysis (MS2) based on the intensity data collected in the first MS1 scan [1]. DIA instead cycles through the entire mass-to-charge ratio (m/z) range of the isolation window without considering any information about the precursors [2]. DIA leads to more complex data because fragmented peptides are not directly tied to a particular precursor and many precursors may be fragmented simultaneously [2].

One method of improving DIA data analysis is to add additional dimensions of separation that make it easier to distinguish similar peptides. Retention time and ion mobility can both be used as dimensions of separation for improving the identification of peptides in high throughput mass spectrometry proteomics. The number of peptides that can be resolved from each other is the peak capacity and, although the two dimensions are not fully orthogonal, Meier *et al*. estimate that a ten-fold increase in analytical peak capacity could be achieved with an average ion mobility resolution of 60 [3]. Another reason to add ion mobility as an additional dimension is that it shows less variation across experiments than retention time which is typically sensitive to the exact experimental conditions present during separation [3]. This means that ion mobility may be a more reliable feature for peptide identification across instruments and laboratories.

Liquid chromatography is a method of peptide separation that sorts peptides based on how they interact with two different phases in a chromatography column [4]. The stationary phase is often a packed column of beads, the surface of which may be coated with other compounds. A mobile phase, in this case a liquid containing the peptide mixture is passed over the stationary phase and peptides elute at different rates depending on the extent to which they interact with the two phases. The time that it takes a peptide to elute is the retention time.

Recently, there has been increasing interest in ion mobility that has been driven by improvements in instrumentation and experimental methods that allow for the analysis of complex samples. Ion mobility is a method of separation that uses the size of a peptide as the discriminating factor [5, 6]. Collisional cross section (CCS) refers to the size and shape of peptide ions in the gas phase as a cross-section of the molecule and represents the rotational average of the ion’s gas phase conformation. In both drift tube and the more recent trapped ion mobility spectrometry (TIMS) technique for ion mobility measurement, ion mobility is reported as the reduced ion mobility coefficient *k_0_*, which can be converted to a CCS value using the Mason-Schamp equation (eq. 1) [7, 8].

Recent advances in the technology, specifically TIMS coupled with time-of-flight instruments (TIMS-TOF) and parallel accumulation-serial fragmentation (PASEF), allow for rapid measurements of peptide ion mobility [9]. In the established technique of drift tube ion mobility, a weak electrical field is used to drag ions through an inert gas while, in TIMS, an electrical field holds the ions stationary against the flow of gas and is then lowered to release the ions in a controlled way (fig. 1 B). This process results in the ions being sorted according to their size as larger particles experience greater forces from the gas and ions with higher charges experience greater forces from the electric field. PASEF uses two TIMS devices to first accumulate ions and then transfers them to the second TIMS device which releases them sequentially in reverse order of CCS [9].

**Figure 1:**
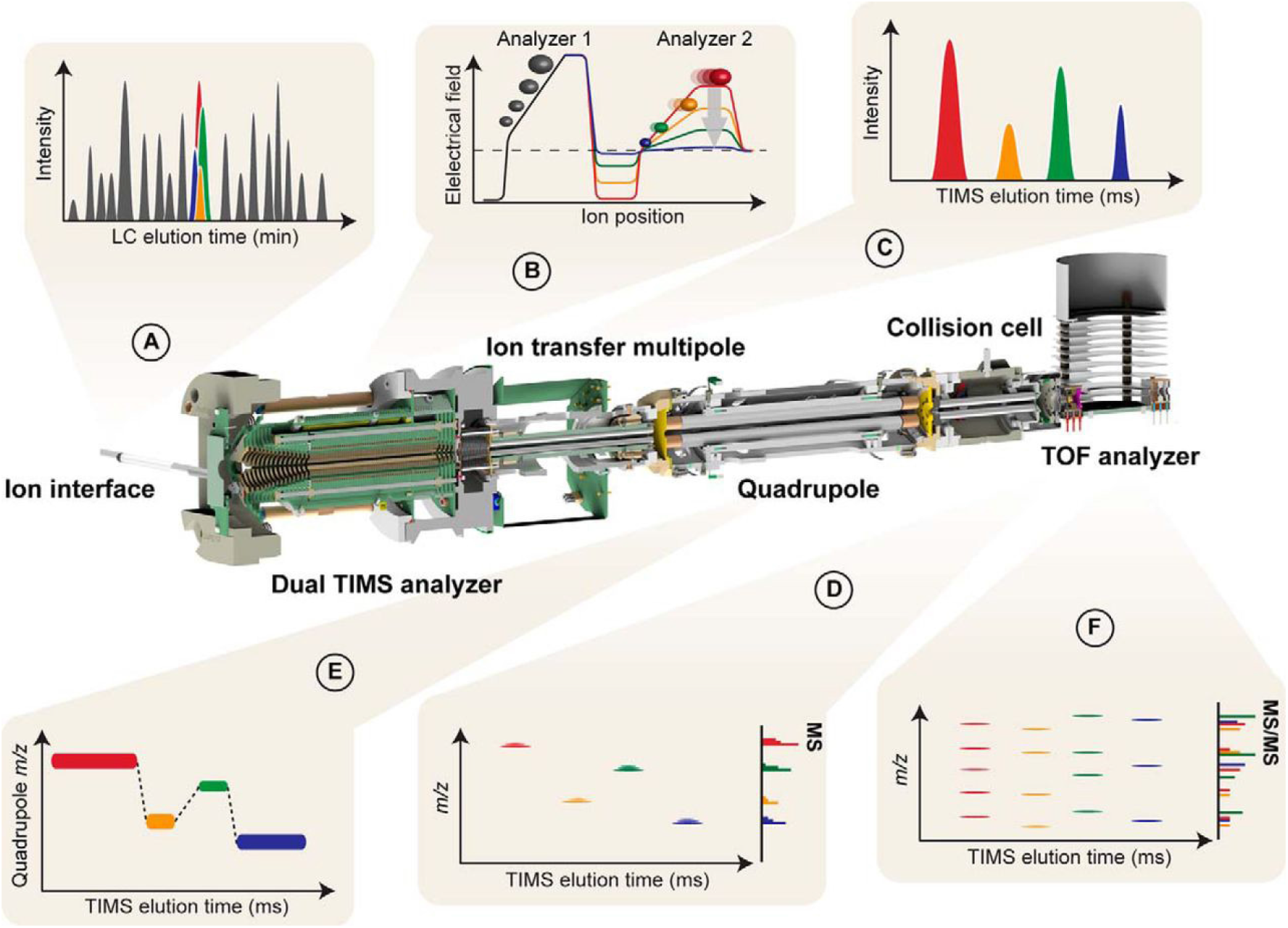
Figure by Meier *et al*. [9] **Online Parallel Accumulation - Serial Fragmentation (PASEF) with the timsTOF Pro.** A, Peptides eluting from the chromatographic column are ionized and enter the mass spectrometer through a glass capillary. B, In the dual TIMS analyzer, the first TIMS section traps and stores ion packets, and the second resolves them by mobility. C, D, Ion mobility separated ions are released sequentially from the second TIMS analyzer as a function of decreasing electrical field strength and yield mobility-resolved mass spectra. E, In PASEF MS/MS scans, the TIMS analyzer and the quadrupole are synchronized and the quadrupole isolation window switches within sub-milliseconds between mobility resolved precursor ions of different m/z. F, This yields multiple ion mobility resolved MS/MS spectra from a single TIMS scan, and ensures that multiple trapped precursor ion species are used for fragmentation. Non mobility-resolved MS and MS/MS spectra are projected onto the right axes in (E) and (F) for comparison. [9]

DIA data analysis relies on DDA libraries to match MS2 spectra and identify peptides [2]. DDA experiments are run to generate libraries containing experimental values for peptide retention times, m/z, relative intensities, and precursor ion mobilities. These libraries are often calibrated for a specific MS instrument making it difficult for different laboratories to share experimental libraries [10]. Additionally, existing libraries may not contain ion mobility measurements since the labs performing the experiments to generate libraries have not had access to instruments capable of measuring ion mobility. Predicted library values promise to eliminate the cost of acquiring new experimental libraries, increase reproducibility, and allow for the augmentation of existing libraries with new attributes.

There has been a trend toward the application of deep learning to predict spectral library attributes with much of the work focused on MS2 spectra and retention time. Much less time has been dedicated to ion mobility prediction despite its importance as an additional dimension of separation in increasingly complex LC-MS-MS data. Recently Meier *et al*. reported a 1.4% median relative error for CCS prediction using an LSTM model [3].

Early methods of prediction use simple neural networks and manually engineered features derived from the amino acid sequence and physio-chemical properties [11]. However, it is unclear what features should be used and how they should be assigned importance. It would be impractical to consider all possible derived features. Deep learning models address this problem by learning both the features and their importance without the need to manually annotate or determine peptide properties. Additionally, deep learning has lead to advances in both genomics [12] and in proteomics, for the prediction of fragmentation spectra and retention time [13, 14, 15, 16].

Two of the most popular models for natural language processing (NLP) tasks are long shortterm memory (LSTM) [17] networks and convolutional neural networks (CNN). Both of these networks can take sequence data as inputs, but while the LSTM processes input sequentially, the CNN can process the entire input simultaneously through parallel operations. In general, LSTMs attempt to understand input by remembering previous inputs and using them as context for the current input, thus allowing them to apply both local and distant information to find features of the input [17]. CNNs examine local sections of the input to find features and then build more complex and abstract features from the features determined in the previous layer [18]. Due to the differences in how input is processed, LSTMs may have an easier time using longer range context information while CNNs should be able to better leverage local information. Both types of information may be important for determining peptide collisional cross section because interactions between close amino acids or those at a distance could affect the conformation of the peptide.

A new type of model, the transformer, has performed very well for NLP tasks such as machine translation. The core mechanism of the transformer architecture is self-attention which is a mechanism of scoring each word in the input with every other word in the input based on relatedness. The benefits of using a transformer compared to LSTMs and CNNs are their ability to learn about both long and short term interactions [19]. For the task of predicting CCS it is unknown if this will provide any benefit as most peptides are relatively short (less than 40 amino acids) compared to the length of a piece of text that may be used in a translation task.

In this project, we set out to compare retention time and CCS predictions on three different machine learning architectures: LSTM, CNN, and transformer. The purpose of each model is to take in a peptide sequence and produce a scalar output of either retention time or collisional cross section (fig. 2). We present an objective comparison of the three model types trained and tested on the same data set and show that the LSTM and CNN architectures achieve similar performance while the transformer has much lower performance.

**Figure 2:**
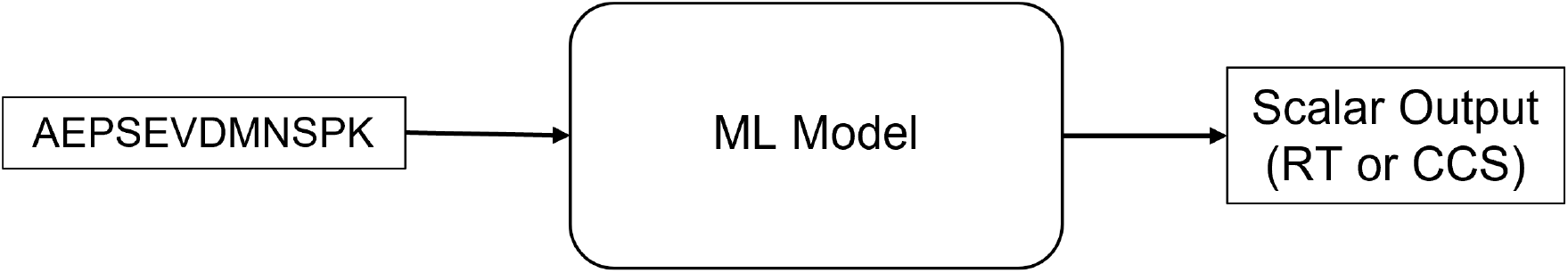
The overall goal of this work is to build machine learning models capable of taking in a peptide sequence and producing a scalar outut of either retention time or collisional cross section

## 3 Methods

### 3.1 Data

The data used in this study was obtained by Meier *et al*. using TIMS-TOF PASEF (parallel accumulation-serial fragmentation) on proteome digests of five organisms and was downloaded from the PRIDE repository with the data set identifier [PXD19086](https://www.ebi.ac.uk/pride/archive/projects/PXD019086) [3]. This data set includes 559979 peptides with sequence lengths ranging from 7 to 55 amino acids and charges fro 2+ to 4+.

Since mass spectrometry experiments report ion mobility as 1/k0 where k0 is the reduced ion mobility, Meier *et al*. used the Mason-Schamp equation 1 [7, 8] to convert 1/k0 to CCS values. This equation is also used to convert predicted CCS values back to 1/k0 in order to predict values for existing libraries.

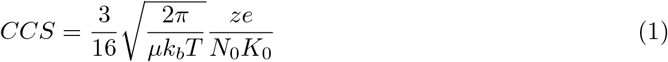

For retention time prediction, only the largest of the experiments, HeLa_Trp_2, was used from the Meier *et al*. data set because retention time is known to have significant variations between experiments. There were 120119 peptides from this experiment.

In all cases, when data was split into train, test, and validation data sets, we ensured that no sequences were present in more than one data set. This was especially important in the case of modified sequences because including the same sequence with different modifications in different data sets could have caused data leakage and lead to artificially good results.

### 3.2 Peptide Encoding

Each model uses the built-in PyTorch embedding layer. From the modified peptide sequence, modification notations were replaced with single letter codes. The modifications present in the data set are oxidation of methionine, acetylation, and carbamidomethylation of cysteine, which were represented as m, c, and a respectively. The single letter codes are then converted to integer representations using a simple dictionary and then to a corresponding embedded vector.

Initially, the peptide charge was appended to the output of the encoder portion of the model before being passed to the fully connected network; however, there was significant improvement in the median relative error of the transformer when the charge was available as part of the model input while not significantly diminishing the performance of the CNN or LSTM models (fig. 3). The peptide charge was provided globally as part of the input by creating a vocabulary dictionary where the keys were the amino acid codes appended to charge (ie. A4 instead of A as done previously). This means that the charge information was available with every token in the input sequence which may have been useful because the position of each charge is not known and could provide important context information.

**Figure 3:**
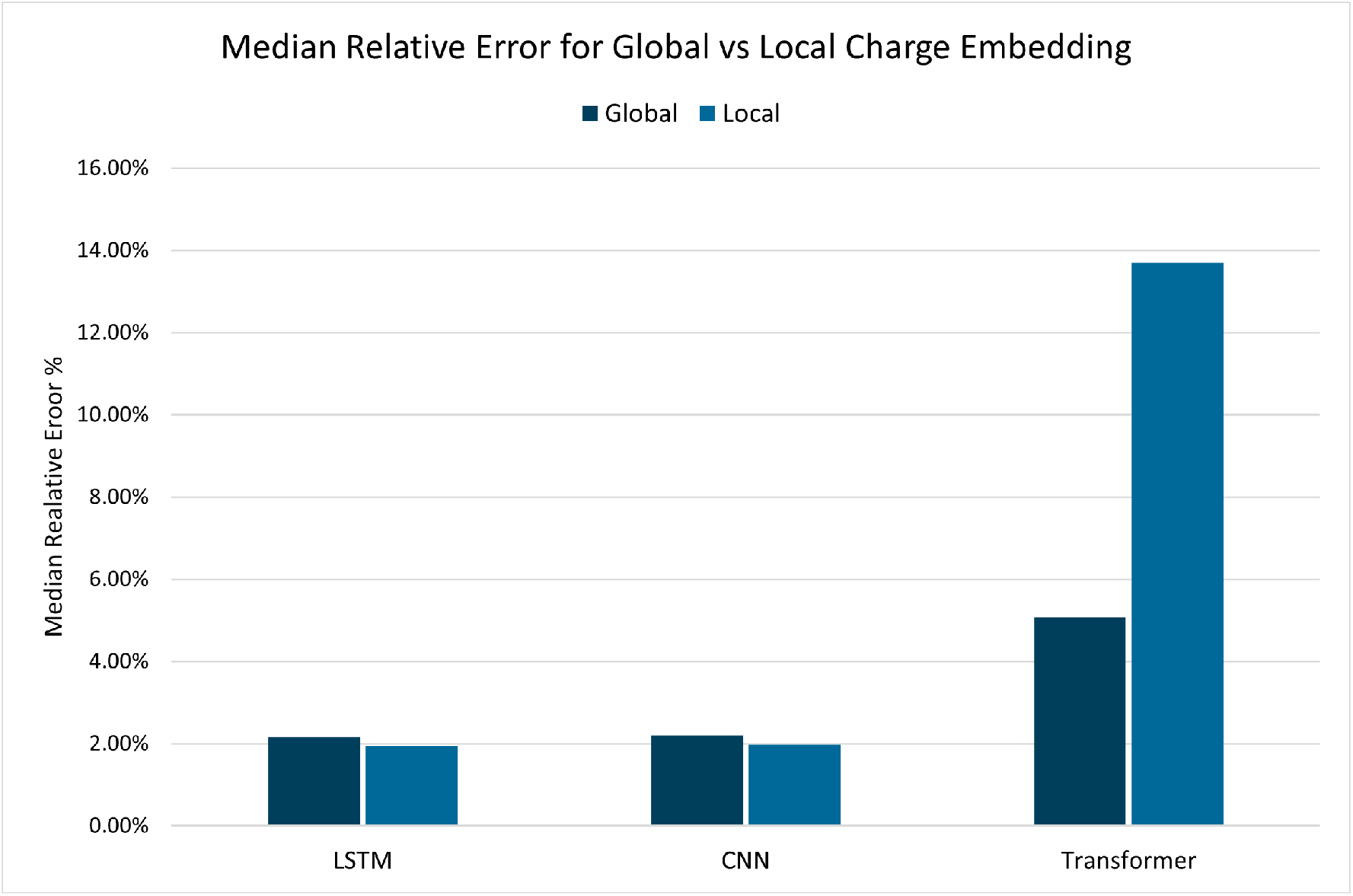
Global vs Local Charge for CCS prediction: When switching from appending the charge to the end of the encoded sequence (local) to incorporating charge information into the model input (global) we can see somewhat diminished performance in the LSTM and CNN, but significant improvement in the transformer. The LSTM performance gets worse with an increase from 1.94% to 2.16% error. Similarly, the CNN performance moves from 1.96% to 2.2% error. The transformer sees significant gains with performance improving from 13.7% to 5.07% error.

**Figure 4:**
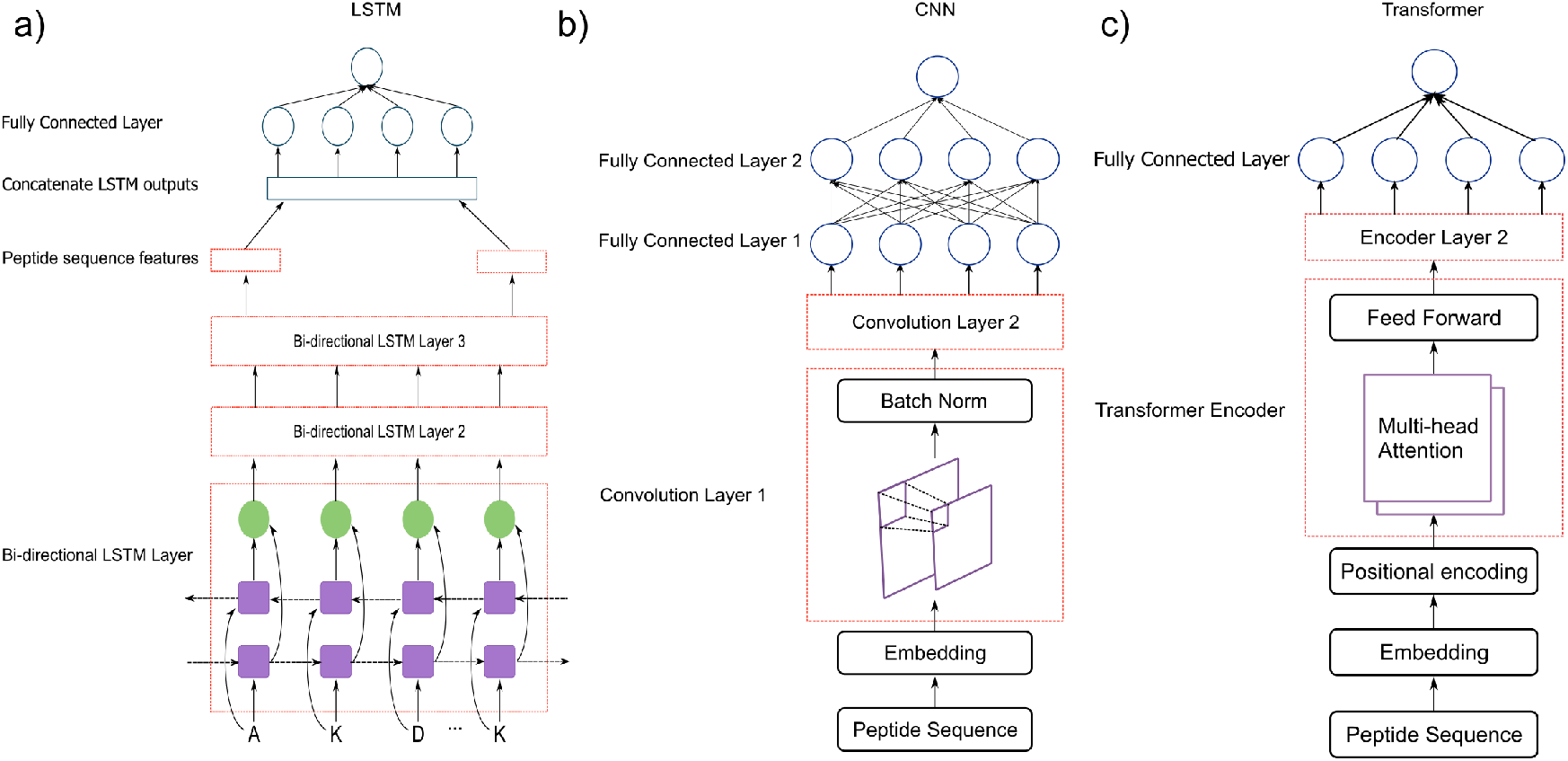
Model Architectures For CCS Prediction: **a)** LSTM architecture consisting of three bi-directional LSTM layers and a two layer fully connected network. **b)** CNN architecture with two convolution layers where each layer is made up of convolution and batch norm sub-layers. This is followed by a three layer fully connected network. **c)** The transformer architecture with an encoder consisting of two layers, each with two attention heads followed by a feed forward network, followed by a two layer fully connected network.

### 3.3 Models

#### LSTM

Recurrent neural networks (RNNs) are a type of network where the input to each self-connected linear unit consists of the input at the current time-step and information on what has been seen in previous time-steps. In these networks, short-term memory refers to the activations of feedback connections that store information about recent inputs and long-term memory refers to the changing of weights over the course of many time-steps in the input [17]. RNNs suffer from the vanishing gradient problem, where error gradients get very small as they are backpropagated causing the network to forget information from much earlier in the sequence. Long short-term memory addresses this problem using multiplicative input and output gates that protect the unit from learning from irrelevant inputs and other units from irrelevant outputs respectively [17].

This model consisted of three bi-directional LSTM layers with 128 units each. Bi-directional layers are used here because the sequence of amino acids is not strictly directional and there may be important contextual information to be gained by considering the sequence both forward and backward. After passing through the LSTM layers, the hidden state of the last time-step for each of the forward and backward directions are concatenated and then input into the fully connected network. Two fully connected layers are used to make the single numerical CCS prediction. The first of the two layers uses a ReLU activation function.

#### CNN

CNNs are common deep learning architectures for image recognition and NLP due to their ability to detect local features and then combine them to form larger features [18]. This means that CNNs should easily recognize the local features of peptides (short-range interactions), but may do poorly if longer range interactions, where an amino acid interacts with one that is farther away, are more important. CNNs work by using a kernel, also called a filter, that is a matrix of values to perform an element-wise dot product on a section of the input that reduces the section to a single feature. The kernel slides across the input data moving by a number of rows or columns at a time known as the stride. In the second convolution layer, the convolution is performed on the features determined from the first step, thus allowing the network to build larger features from smaller ones.

The CNN used here has two convolution layers each followed by a batch normalization layer. The kernel size used in each convolution had dimensions of 9 and the embedding dimension, which means that nine amino acids (or less, if the sequence is shorter) are examined in each convolution, and the stride is one. The output of the convolution layers is then passed to a fully connected network with three layers.

#### Transformer

The basic transformer architecture consists of stacks of encoders and decoders where each is comprised of a self-attention and point-wise feed-forward layer [19]. Before going to the encoder, inputs are first passed through a positional encoding layer to capture information about the relative order of the tokens in the input sequence. The self-attention mechanism, know as “Scaled Dot-Product Attention”, works by calculating the output vector as a weighted sum of a value vector where the weights are derived from the similarity of key and query vectors as shown in equation 2 [19].

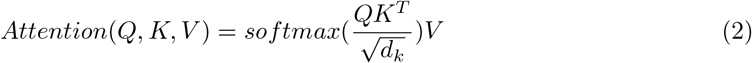

Since the goal here was to predict a single output value, a simple linear decoder is used. This decoder is a two layer fully connected network with a ReLU activation function on the first layer. We also use the built-in PyTorch embedding and a positional encoding prior to the PyTorch TransformerEncoderLayer component. Two transformer layers and two attention heads were used build the encoder block.

#### Retention Time Prediction

The models use slightly different hyper-parameters for retention time prediction that were determined using a hyper-parameter sweep, as was done for CCS prediction, that tests a range of values for each model hyper-parameter. For each model, some differences from above are that the embedding dimension was 64 and the batch size was 4. A smaller batch size may be better here because the training data set is much smaller and so the models have more opportunities to update their parameters before they start seeing the same data again. It is unclear why a smaller embedding dimension is preferred for retention time prediction. It may be that the exact size of the embedding dimension is not that important, which is supported by the fact that we saw similar but somewhat worse performance with an embedding dimension of 128, but this was not investigated further. Additionally, the transformer had 8 layers and 8 attention heads, rather than the 2 as described for CCS prediction.

### 3.4 Training

All models were trained on a NVIDIA GeForce GTX 1650 SUPER graphics card for 20 epochs with a batch size of 64. The approximate training times were 1.5 hours for the LSTM, 45 minutes for the CNN, and 30 minutes for the transformer with some variation between runs. All models were trained using mean squared error as the loss function, but here we report the median relative error as it is better for making comparisons between models.

Models were trained using the PyTorch implementation of the Adam optimizer with a starting learning rate of 0.001. The learning rate is reduced by a factor of 0.1 after the first training epoch and after that by a factor of 0.1 by a learning rate scheduler that reduced the learning rate when the median relative error stopped improving. For all other parameters, the PyTorch defaults were used. The hyper-parameters used, such as the dimensions of fully connected layers and the number of layers were determined using a hyper-parameter search.

The software Weights and Biases was used for experiment tracking throughout the process of developing these models [20].

## 4 Results

### 4.1 Retention Time

Comparing the three model architectures, we can see from figure 5 that the LSTM and CNN showed similar performance while the transformer performed somewhat worse. Although each model had a similar median relative error, the transformer has a much larger distribution of error. For each of the histograms in figure 5 the x-axis is confined to the range [−50, 50] so that the distribution is clear. Outside of this range the LSTM, CNN, and transformer had 152, 173, and 887 values respectively. Most of the distribution is contained between 0 and −20% for the LSTM and CNN and between −20 and 20% for the transformer. This means that while all three models have similar median errors, the transformer was less accurate. The plot of experimental vs predicted values for the transformer (fig. 8 f) also shows less accuracy with a Pearson correlation of 0.938 compared to 0.983 and 0.979 for the LSTM and CNN. We can also see that all three models tend to underestimate the retention time, as shown by more of the errors being below zero, however this effect is more pronounced in the LSTM and CNN. This is a trend that continues for CCS prediction (fig. 8).

**Figure 5:**
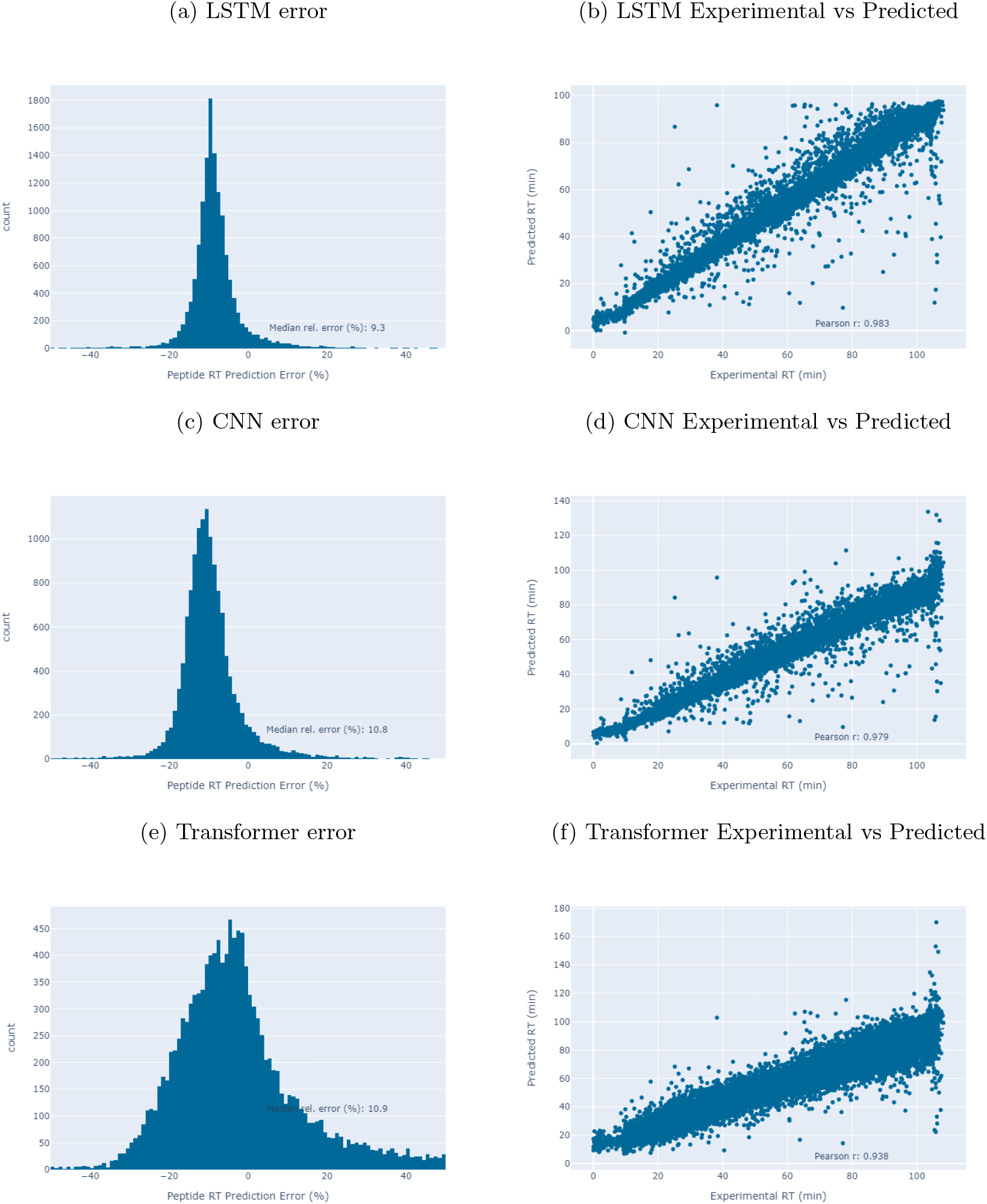
Retention Time Prediction. a) Distribution of the percent error for the LSTM model. b) Experimental vs Predicted RT values for the LSTM. c) Distribution of the percent error for the CNN model. d) Experimental vs Predicted RT values for the CNN. e) Distribution of the percent error for the transformer model. f) Experimental vs Predicted RT values for the transformer.

The models presented here were unable to perform as well as DeepRT [21] on retention time prediction. Since DeepRT was shown previously to outperform an optimized LSTM, it was expected that it would outperform the LSTM and the basic CNN used here. The transformer performed worse than expected but may require additional optimization or data. Figure 9 shows the relationship between data set size and the median relation error for ion mobility prediction and we can see that the transformer shows significant improvement when it is trained on more data.

### 4.2 CCS

The distribution of error was much broader for the transformer than the LSTM or CNN (fig. 8). Most of the error for values predicted by the LSTM or CNN models is between −5% and 5%, while the errors for the transformer are mostly between −10% and 10% with a much more even spread across this range. The LSTM and CNN plots both show spikes just below zero and the number of predictions at each 1% change in error drops rapidly. From these histograms we can also see that the LSTM and CNN models tend to underestimate the CCS values with most of the errors being negative. The transformer does not seem to have this same tendency and shows a roughly even split of errors above and below zero. We also see that the LSTM and CNN both achieve median relative errors of 2.1% and the transformer error is 4.6%.

We also see very similar performance of the LSTM and CNN in the experimental vs predicted CCS plots in figure 8. Both the LSTM and CNN show very high correlation with Pearson correlations of 0.992 and 0.994 respectively. The transformer has a Pearson correlation of 0.976 and we can see that it predicts values farther from experimental at higher CCS values. These higher CCS values correspond to higher charge states on which all models performed worse, especially the transformer (fig.7).

From figure 7, we can see that LSTM and CNN models had similar predictive capabilities for each charge state, while the transformer performed worse than the other models on all charge states. All models had lower performance with increasing charge state. This is likely due to the decreasing number of peptides with each charge state in the training data. From figure 6, which shows the number of peptides with each charge state in the training data, we can see that there are less than half as many peptides with a charge state of 3+ compared to peptides with a charge state of 2+, and less than one tenth as many peptides with a charge of 4+ compared to 2+. With far fewer examples of peptides with higher charge states, it is not surprising that the model was not able to learn as well.

**Figure 6:**
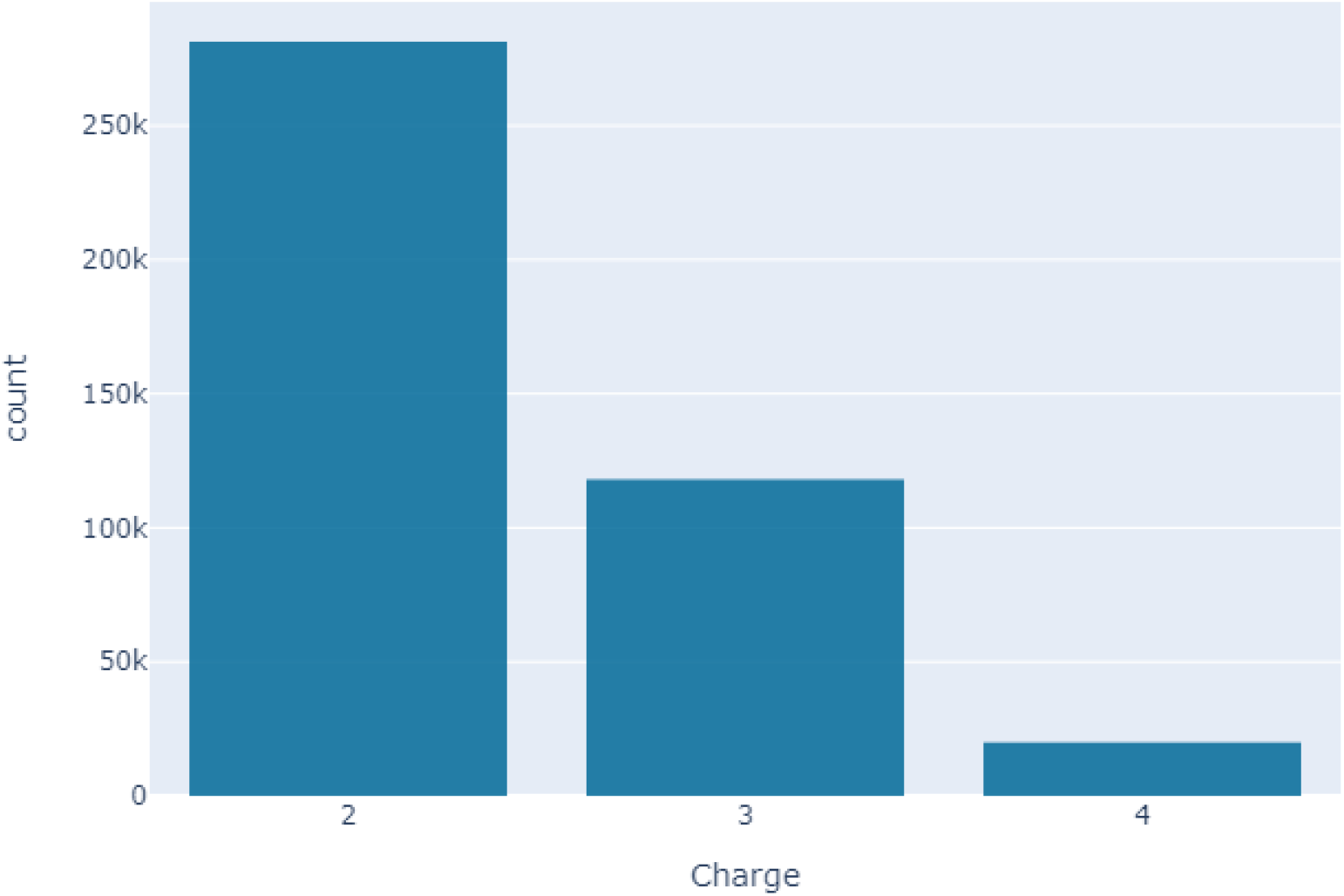
Distribution of charge states in the training data. As charge increases, the number of peptides decreases rapidly. There are 281,328 peptides with a charge of 2+, 118,114 peptides with a charge of 3+, and 20,013 peptides with a charge of 4+.

**Figure 7:**
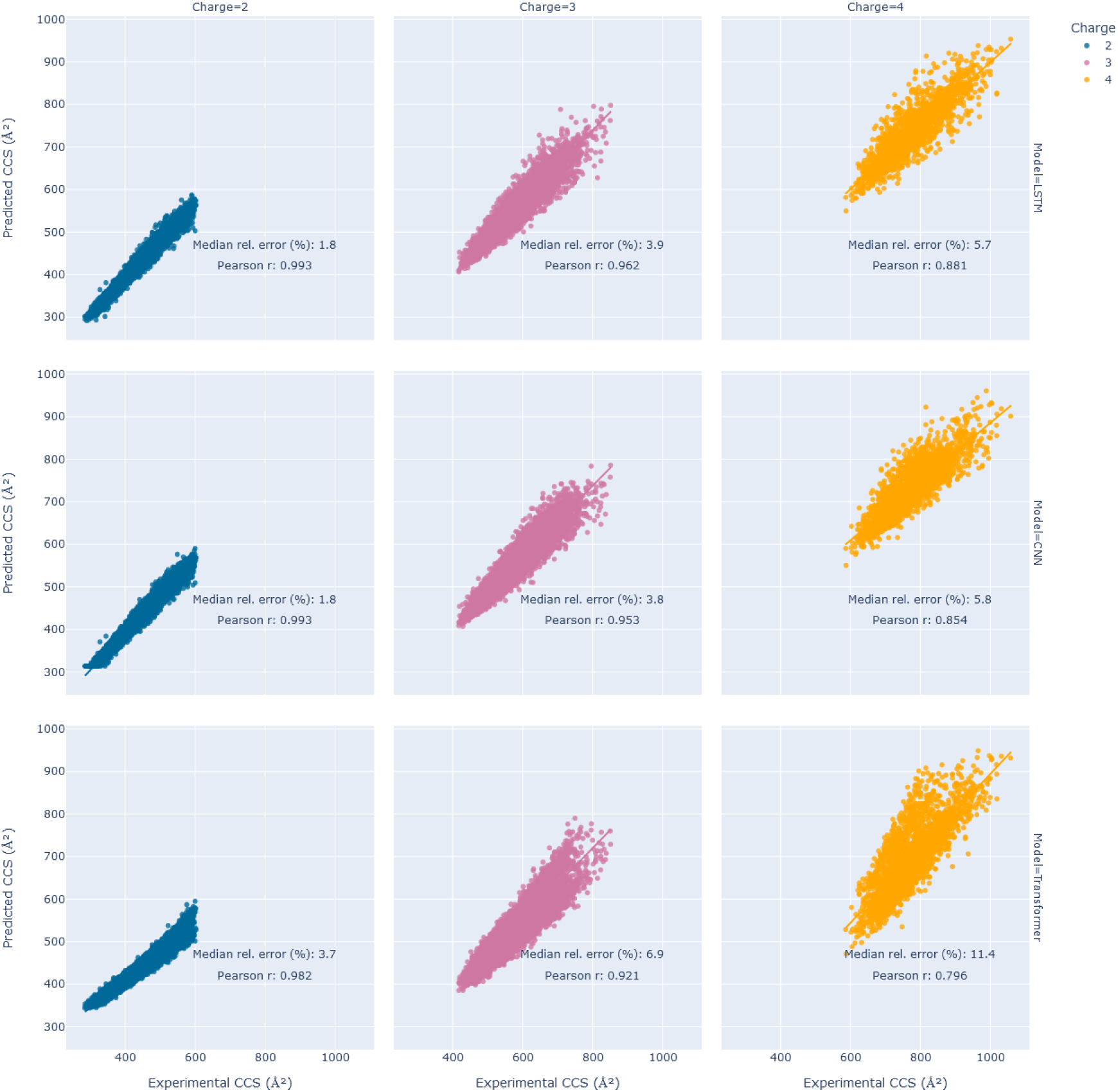
Experimental vs predicted CCS value. Each column shows the experimental vs predicted CCS values for a charge state with the first, second, and third columns showing charge state 2, 3, and 4 respectively. The top row shows the results for the LSTM model, the middle row for the CNN, and the bottom row contains results for the transformer.

**Figure 8:**
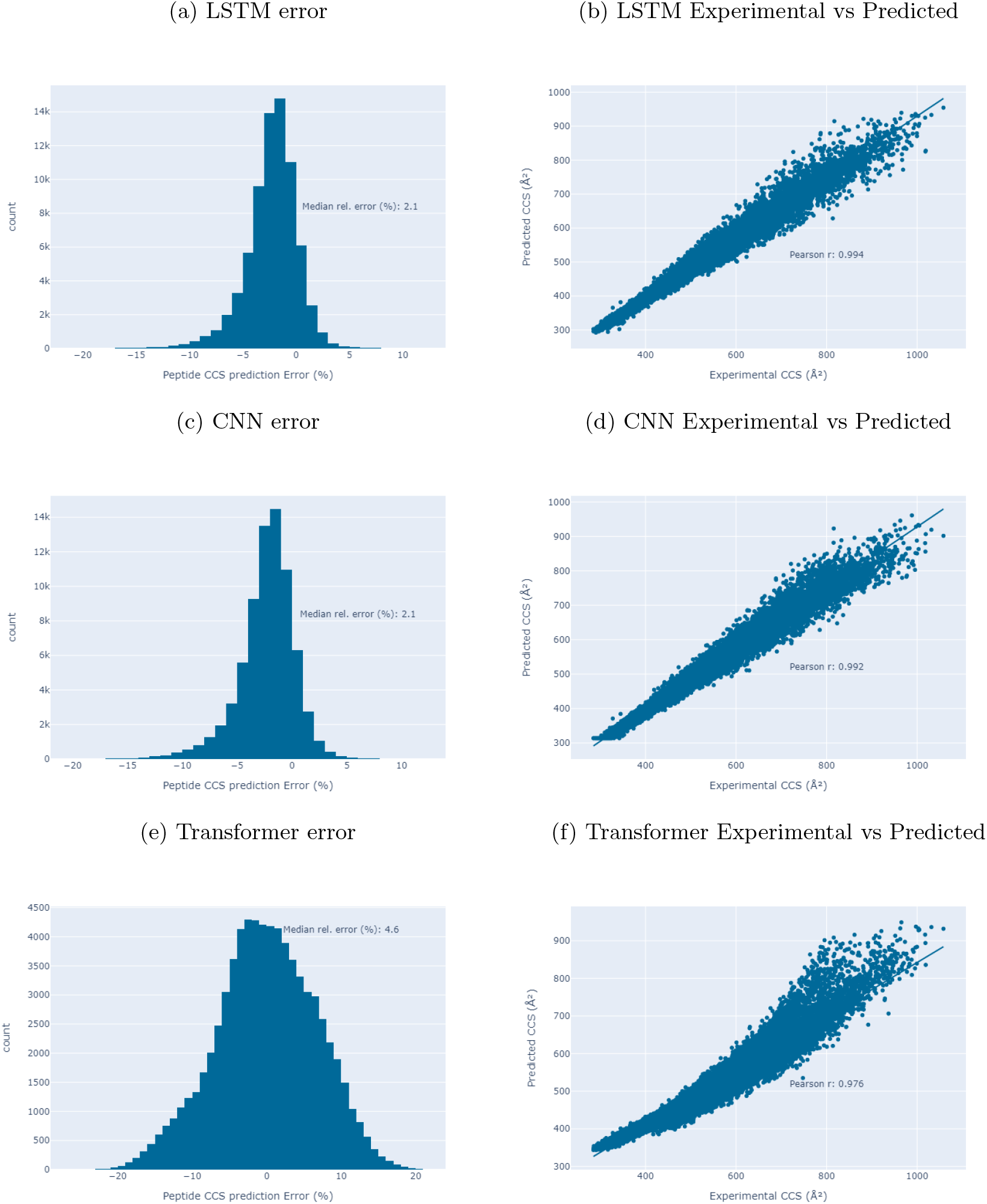
CCS Prediction a) Distribution of the percent error for the LSTM model. b) Experimental vs Predicted CCS values for the LSTM. c) Distribution of the percent error for the CNN model. d) Experimental vs Predicted CCS values for the CNN. e) Distribution of the percent error for the transformer model. f) Experimental vs Predicted CCS values for the transformer.

Explaining the poorer performance of the models at higher charge states as being due to a lack of training examples relies on the assumption that the charge is actually and important piece of information in making these predictions. Although the present work does not explore the interpretability of the models used, charge does seem to be important for leaning to predict CCS, especially for the transformer, as shown in figure 3 where there was a significant improvement in performance when the charge information was made more available to the model.

## 5 Discussion

Here we have compared the performance of three common deep learning architectures on the tasks of predicting the scalar properties of peptide RT and CCS. As shown in figure 7, the LSTM and CNN models perform very similarly overall and for different charge states while the transformer lagged behind in performance both overall and for each charge state. We suggest that the poor performance of the transformer and the reason why that performance degrades more quickly with increasing charge state than in the other models is due to the much smaller sample size of peptides with higher charges.

The models presented here are consistent with their performance of recently developed models for retention time and collisional cross section prediction, but they do not match the performance. We have seen that the LSTM and CNN perform similarly, but would like to bring the performance of the transformer closer to that of the other models. It may be possible to do this by increasing the sample size, increasing the size of the model, or augmenting the architecture. From figure 9, we can see that transformer is the most affected by sample size and so may benefit from a larger data set, although, this is unlikely to completely close the gap between the transformer and the other models. Additionally, modifications to the relatively basic model architectures used here could yield significant improvements. One architecture to explore in future work is the conformer [22]. The conformer combines the CNN and transformer architectures to make use of the strengths of each. The transformer is better able to leverage global interactions while the CNN is better at using local features.

**Figure 9:**
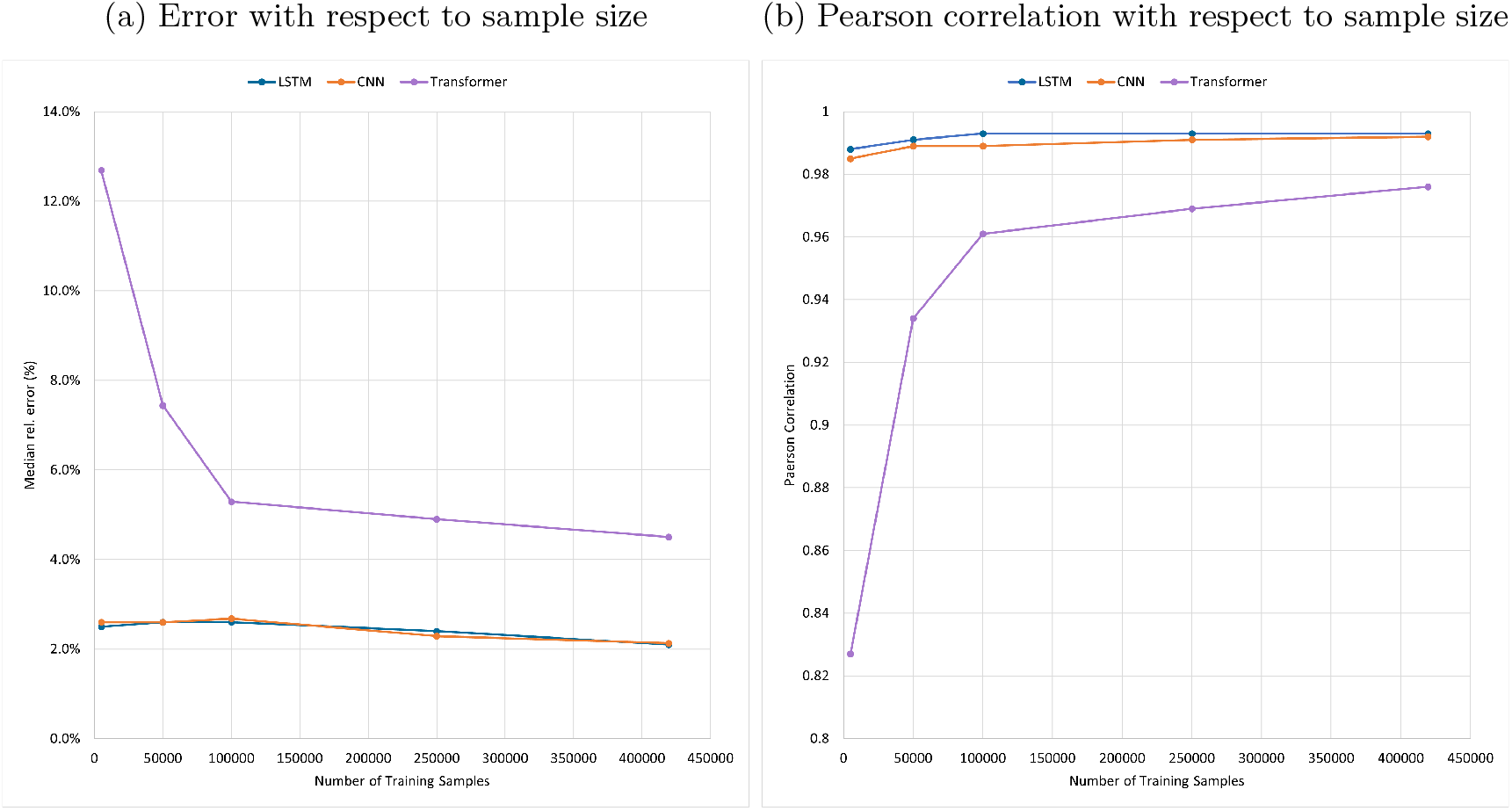
**a)** The relationship between median relative error and training data set size. **b)** Pearson correlation coefficient with respect to the training data set size.

Another possible explanation for the lower transformer performance is that it has far fewer learn-able parameters. The LSTM, as described in the methods for CCS prediction, has just over a million learn-able parameters and the CNN has twice as many at just over two million. The transformer, however, has only about 250,000 learn-able parameters. It is possible that the transformer may benefit from increasing the size of the model and the number of learn-able parameters as it may increase the model’s capacity to learn. Two ways of doing this are to increase the number of encoder layers and increase the number of attention heads. A similar number of parameters are present in each of the retention time prediction models. Increasing the size of the transformer for better performance is supported by the retention time prediction results because, when determining the hyper parameters to use, a larger number of attention heads and layers lead to better performance on validation data. This was not investigated in detail and the CCS prediction models had previously only been tested with 2 and 4 attention heads/number of layers with no improvement observed.

One reason for trying to keep the architectures as simple as possible is to prevent overfitting and preserve interpretability. The more complex a model becomes, the more parameters it has available to learn. This can lead to models that learn the training data very well, but fit the data so closely that they no longer generalize well to new data. Keeping models simple helps to limit overfitting, but there are also many other options for reducing overfitting such as using dropout or a penalty on the weights. The number of parameters for each model

## 6 Code Availability

The code is available on [GitHub](https://github.com/E-Franklin/DeepLearningForIonMobility).

## 7 Acknowledgements

I would like to thank my supervisor Hannes Röst for guidance in completing this project, as well as all the members of the Röst lab for feedback on this work. The experiments were made possible by a compute allocation from Compute Canada. I would also like to thank the Department of Computer Science and NSERC CGS-M for their financial support throughout the program.

